# Construction of functional biliary epithelial branched networks with predefined geometry using digital light stereolithography

**DOI:** 10.1101/2021.07.19.452904

**Authors:** Elsa Mazari-Arrighi, Dmitry Ayollo, Wissam Farhat, Auriane Marret, Emilie Gontran, Pascale Dupuis-Williams, Jerome Larghero, Francois Chatelain, Alexandra Fuchs

**Affiliations:** Université de Paris, Inserm, U976 HIPI, F-75006, Paris, France; AP-HP, Hôpital Saint-Louis, 1 avenue Vellefaux F-75010, Paris, France; CEA, IRIG, F-38000, Grenoble, France; INSERM U-1279, Gustave Roussy, Villejuif, F-94805, France; Université Paris-Saclay, Inserm, Physiopathogenèse et traitement des maladies du foie, F-94800, Villejuif, France; ESPCI Paris, Université PSL, F-75005 Paris, France

**Author notes:** Corresponding author: Alexandra Fuchs, Address: Inserm U976, porte 15B, Hôpital Saint- Louis, 1 Avenue Claude Vellefaux, Paris, 75010, France.

**Keywords:** Tissue engineering, bile ducts, biliary tree, 3D bioprinting, DLP stereolithography, cholangiocytes

## Abstract

Cholangiocytes, biliary epithelial cells, are known to spontaneously self-organize into spherical cysts with a central lumen. In this work, we explore a promising biocompatible stereolithographic approach to encapsulate cholangiocytes into geometrically-controlled 3D hydrogel structures to guide them towards the formation of branched tubular networks. We demonstrate that within the appropriate mix of hydrogels, normal rat cholangiocytes can proliferate, migrate and organize into branched tubular structures, form walls consisting of a cell monolayer, transport fluorescent dyes into the luminal space and show markers of epithelial maturation such as primary cilia. The resulting structures have dimensions typically found in the intralobular and intrahepatic bile ducts and are stable for weeks, without any requirement of bulk supporting material, thereby offering total access to the basal side of these biliary epithelial constructs.

## Introduction

Liver tissue engineering is an actively developing research area in the field of drug screening to develop more physiologically relevant toxicity assays and models of liver diseases. In addition, it is a promising solution for regenerative medicine to alleviate the need for donor liver tissue. In recent years, significant progress has been made in culturing primary or iPSC-derived hepatocytes in the form of spheroids or organoids to achieve higher differentiation and maintain higher functionality *in vitro*^1–6^. To date, the most advanced example of this approach was described by Takebe et al, who managed to produce liver buds composed of iPS-derived endoderm, endothelial and mesenchymal cells grown to form organoids that were shown to rescue hepatic functions in a mouse model of acute liver failure^7^. Such an approach has the potential to create relevant models of metabolic liver diseases, as well as to ultimately treat the partial failure of liver functions. However, they do not take into account the critical role played by the bile ducts and ductule network in collecting and clearing toxic compounds and metabolites from the hepatic tissue.

The human biliary tree consists of the intrahepatic and extrahepatic sections. The intrahepatic biliary tree is composed of ductules that collect and modify the bile from the bile canaliculi formed by hepatocytes in the liver lobules. The intrahepatic ductules gradually merge into the common hepatic duct which leads to the extrahepatic biliary tree ^8,9^. Over the past few years, there have been a few successful attempts to produce bio-artificial bile ducts (BABD). For example, in 2019, Tysoe et al. have described a way to build a bile duct with a lumen diameter of 250 µm using a rolled polyglycolic acid or a collagen hydrogel layer as a scaffold for cholangiocytes isolated from murine liver and amplified as organoids^10^. Another approach was described by Chen et al. in 2018 in which authors used cholangiocyte-like cells from the murine liver to seed the outer surface of polyethersulfone hollow fiber membranes functionalized with L-Dopa and collagen^11^. The result was a BABD with an inside-out inverted polarity and a diameter of 1 mm. While the above-mentioned works mark obvious progress in extrahepatic bile duct reconstruction, reproduction of the intrahepatic part of the biliary tree is more challenging due to both its size and its complexity. The intrahepatic biliary tree permeates every level of liver anatomy from the hepatic hilum to the liver lobule, with internal luminal diameters ranging from 800 µm in the interlobular ducts down to 10 µm inside the lobules.

Using the relatively popular top-down approach of decellularization and subsequent recellularization of the whole organ, the liver vasculature has been successfully repopulated with endothelial cells^12^. However, the recellularization approach for the biliary tree showed limited success^13^. This may be due to several factors ranging from the intricate biology of cholangiocytes to the tortuous and complex architecture of the intrahepatic biliary tree.

Bottom-up approaches to *de novo* engineering of the intrahepatic biliary tree have been pursued using hydrogel matrices. *In vitro*, extracellular matrix (ECM) substitutes such as type 1 collagen or Matrigel (an ECM hydrogel produced by the mouse Engelbreth-Holm-Swarm sarcoma cell line) have shown promising results both for cholangiocytes alone or in co-culture with hepatocytes^14,15^. Cholangiocytes, known to have a certain degree of *in vitro* morphogenic capability, spontaneously form cysts when cultured in natural or synthetic hydrogels^11,16^ and cystic organoids obtained from primary cholangiocytes have shown to repair intrahepatic biliary ducts when transplanted in a human liver undergoing *ex vivo* normothermic perfusion^17^. However, elongated duct formation *in vitro* in Matrigel or type 1 collagen is rarely seen^18–24^. Another drawback is the lack of control over direction, length, branching, and diameter of occasionally forming tubular structures.

A promising approach to building tissue constructs with controlled geometry is 3D bioprinting. In this field, Lutolf’s team illustrated how extrusion-printing of a high-density suspension of gut epithelial cells in a Matrigel/collagen hydrogel could lead to self-organization of cells into tubes with a luminal diameter of 100-200 µm^25^. To reach an even higher resolution, in particular, to reach features in the range of 10-100 µm, digital light processing (DLP) stereolithography is particularly efficient. This approach was illustrated in a publication by Ma et al., where different types of cells (iPS-derived hepatic progenitor cells, endothelial cells, and adipose-derived stem cells) were encapsulated in imbricated 3D structures mimicking the hexagonal design of the liver lobule^26^. The resulting construct exhibited enhanced hepatic phenotypes and metabolite activity. However, the reconstruction of the biliary network was not addressed in their work.

In our study, we used normal rat cholangiocytes (NRC) and DLP stereolithography to produce Biliary Structures by PhotoPolymerization (BSPPs) with a controlled geometry. After several days of culture, the self-organization of branched tubular structures guided by the predefined geometry of the hydrogel scaffold leads to the formation of BSPPs. These tubes spanning hundreds of microns had a central closed lumen with walls composed of a monolayer of cells with an epithelium phenotype, including the presence of tight junctions and primary cilia facing the lumen. Our results also demonstrate that these cholangiocyte structures possess functional transport properties of bile ducts.

## Results

### Cholangiocytes embedded in 3D photopolymerized hydrogels of defined geometry can proliferate, migrate, and organize into branched tubular epithelial networks

Cholangiocyte cysts which form spontaneously in hydrogel environments, consist of a central lumen surrounded by monolayer of cells with apicobasal polarization^11,16,18,21,27^. When cultured in ECM derived from the decellularized liver^18^, some cysts randomly adopt an elongated shape. However, their size and shape cannot be controlled, and the system is inappropriate for obtaining reproducible models of functional polarized intra-lobular biliary ducts. In an attempt to guide cholangiocytes’ self-organization towards tubes with predetermined geometry, we used DLP stereolithography to fabricate 3D hydrogel structures of various shapes encapsulating biliary epithelial cells **(Fig. 1)**.

**Figure 1.**
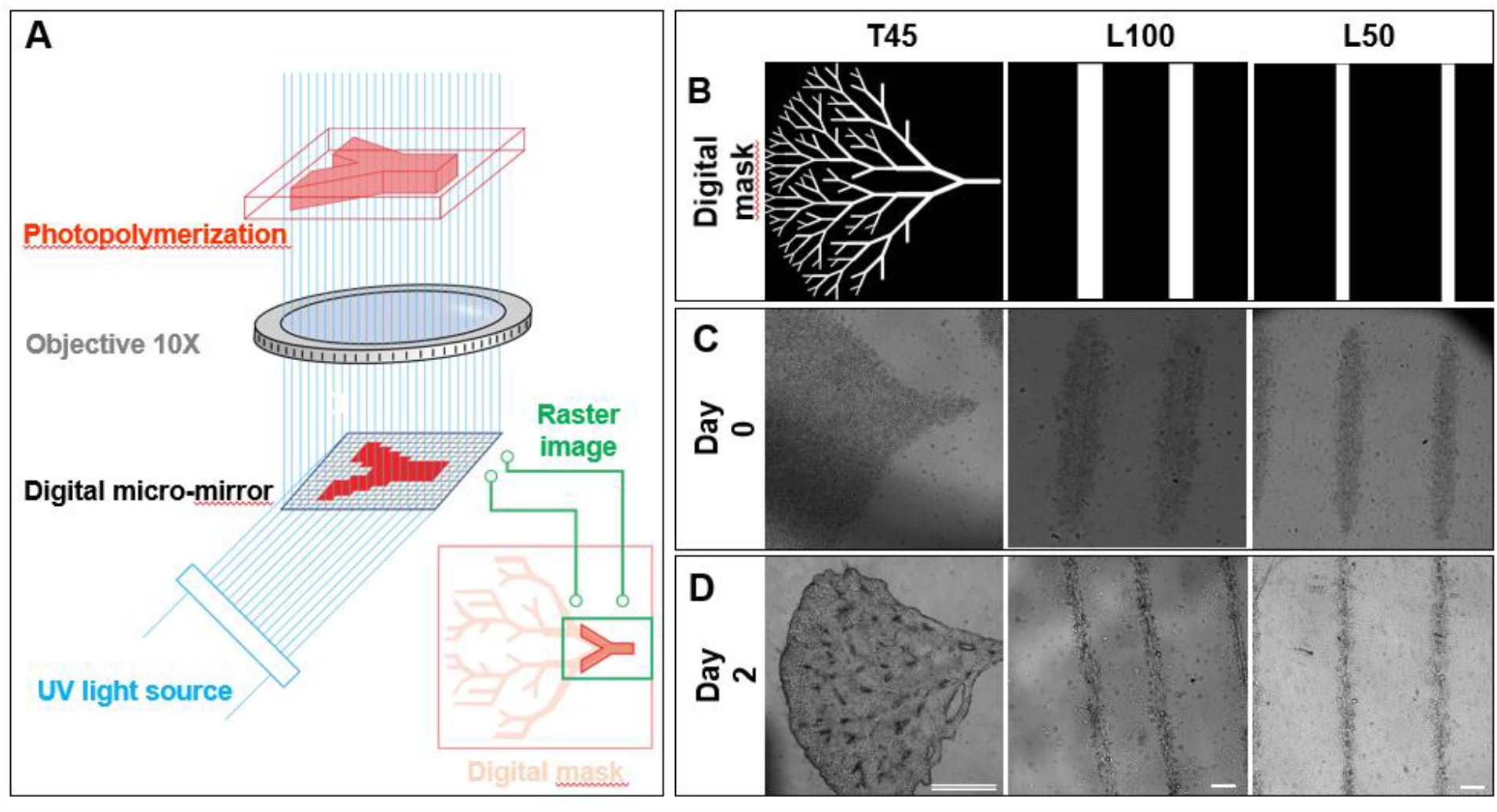
Fabrication of BSPPs. (A) The scheme of DLP stereolithography. (B) Various digital patterns for BSPPs: a tree with 45° branching (T45), and lines of 100 µm (L100) and 50 µm (L50) width. (C, D) Cholangiocytes encapsulated into a correspondingly shaped photopolymerized hydrogel structures just after fabrication (C) and 2 days after the fabrication (D). Double-line scale bar is 1 mm (tree pattern) and single line scale bars are 200 µm (line patterns).

After surveying a panel of ECMs that have been shown to sustain (alone or in a mixture) the culture of epithelial cells^28^, we focused on type I collagen, hyaluronic acid, and fibrinogen. In order to print the selected hydrogels using DLP stereolithography, we screened different formulations of methacrylated collagen (C^MA^), methacrylated hyaluronic acid (HA^MA^) and lithium phenyl-2,4,6 trimethyl-benzoyl phosphinate (LAP) – a cytocompatible photoinitiator^29^, with or without fibrinogen (FG) **(Fig.2)**.

C^MA^ is a photopolymerizable hydrogel produced from collagen that sustains good cell viability following encapsulation and retains natural cell-binding motifs^30,31^. HA^MA^ is also a photopolymerizable hydrogel derived from hyaluronic acid, the latter is found ubiquitously in native tissues and plays an important role in many cellular responses, such as cell signaling or cell proliferation^25–27^. FG is a component of the ECM that mediates cellular functions such as adhesion, spreading, proliferation, and migration of a variety of cell types, including fibroblasts, endothelial and epithelial cells. C^MA^ and HA^MA^ incorporation provides tunability of mechanical properties via variation of the UV dose and material concentration while still maintaining the essential features of the native biliary epithelial microenvironment^32,33^. The incorporation of FG changes the topology of the extracellular matrix in order to provide a surface for cell adhesion, migration, and matrix remodeling^34^. As illustrated in **Fig. 2A**, C^MA^ and HA^MA^ when used alone or in combination with FG could be photopolymerized with suspended cholangiocytes. However, C^MA^, HA^MA^/FG, and C^MA^/FG hydrogels were rapidly degraded in the presence of living cells and thus did not offer enough stability to sustain directed growth of any organized multicellular structure. This degradation was directly linked to the presence of living cells as the same hydrogel structures shown excellent stability over time in the absence of cells. In contrast, when all three components were mixed in a ratio of 0.2% (w/v):0.3% (w/v):0.14% (w/v) C^MA^/HA^MA^/FG, photopolymerized hydrogel structures showed substantially higher stability over time and we observed migration and reorganization of the cholangiocytes within them.

**Figure 2.**
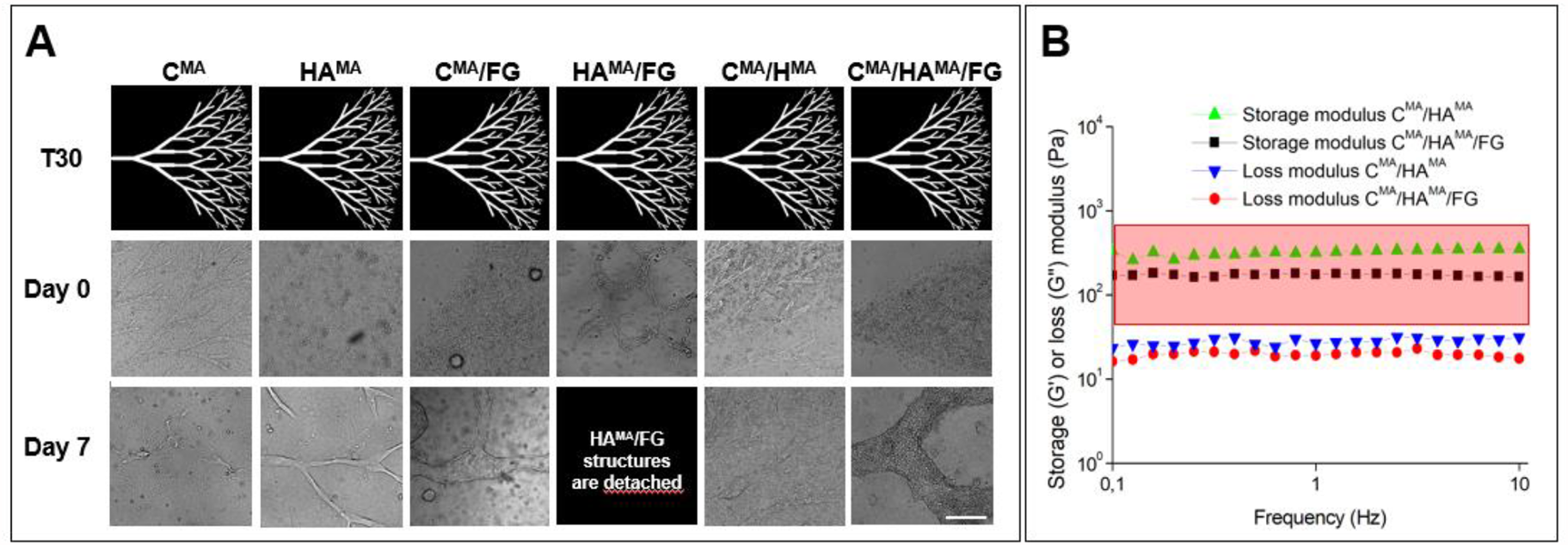
*In vitro* characterization of various hydrogels for BSPPs. (A) Cholangiocytes encapsulated in various photopolymerized 30° tree hydrogels (T30) just after photopolymerization and 7 days after photopolymerization. It is worth noting that FG is not photopolymerizable using the LAP photoinitiator. Scale bar is 200 µm. (B) Rheological analysis of the photopolymerized hydrogels used to encapsulate cholangiocytes in comparison with the storage modulus range for optimal concentration of decellularized liver ECM enabling tubulogenesis according to Lewis et al. (red framed rectangle)^18^.

Storage modulus (G′) and loss modulus (G″) were measured via frequency-dependent oscillatory rheological analysis to investigate the viscoelastic features of the fabricated hydrogels. As depicted in **Fig. 2B**, both CMA/HAMA and CMA/HAMA/FG formulations exhibited a classical hydrogel feature (G′ greater than G″), where G′ and G″ were kept nearly constant in the entire frequency range (0.1–10 Hz). These measurements also showed that the addition of FG in the photopolymerizable hydrogel formulation for BSPPs allowed the formation of a softer matrix compared to C^MA^/HA^MA^. Since matrix stiffness is known to have significant effects on cell behavior^35^, the increase in stiffness for C^MA^/HA^MA^ mix may be responsible for the limited migration that is observed for NRC encapsulated within such photopolymerized structures (**Fig. 2A**, C^MA^**/**HA^MA^ **day 7**). It is interesting to note that the storage modulus of the C^MA^/HA^MA^/FG photopolymerized hydrogel is in the same range of 50-800 Pa as the one measured by Lewis et al. for ECM derived from decellularized livers^18^ in which cholangiocytes could reorganize *in vitro* into cystic and elongated structures. In parallel, by optimizing UV doses and concentration of LAP, we confirmed that cells had good viability 4 hours and 24 hours after printing **(Fig. S1)**.

We also explored various patterns of lines ranging from 10 to 500 µm in width and trees with branching angles of 30, 45, and 60° (**Fig. 1 and Fig. S2**). In all cases, we found that, just after photopolymerization, cells were trapped in an area larger than the one defined by the optical mask (**Fig. 1C**). This is most likely the result of light scattering, thereby inducing partial photopolymerization in adjacent regions and decreasing the expected resolution^36^. Nevertheless, after two days in culture, the structures had condensed (**Fig. 1D**) and the resulting structures were close in size to the original projected pattern, as if the cells had clustered together around the harder polymerized backbone.

**Figure S1.**
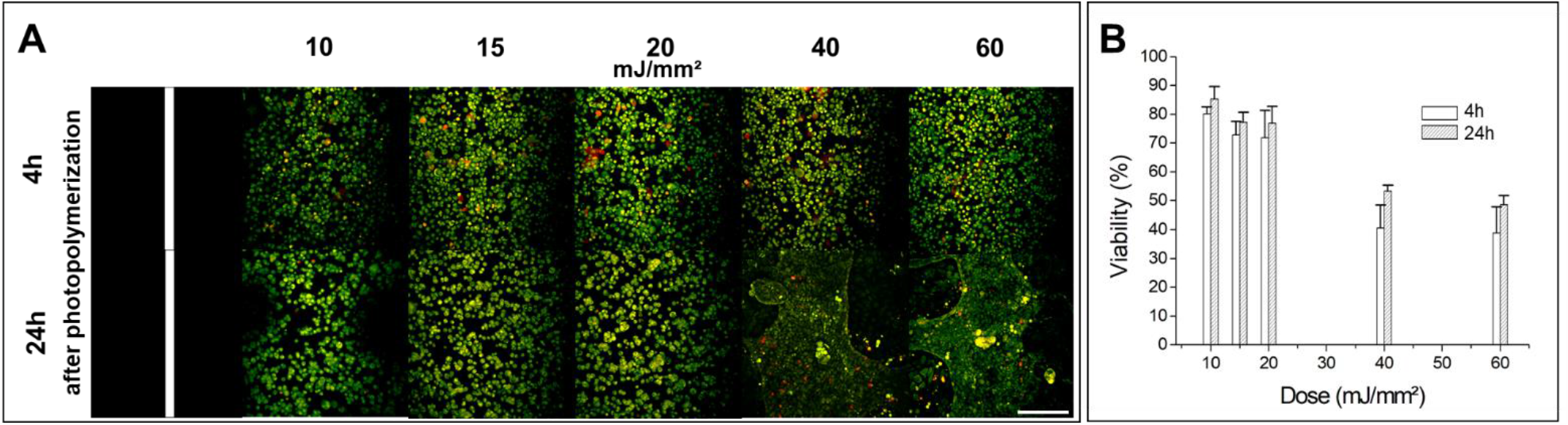
Cell viability analysis in BSPPs. (A) Maximal projection images showing staining of living cells by calcein (green) and of dead cells by ethidium homodimer-1 (red) encapsulated into BSPPs 4h and 24h after photopolymerization. BSPPs were obtained using: 10 mJ/mm^2^, 15 mJ/mm^2^, 20 mJ/mm^2^, 40 mJ/mm^2^, and 60 mJ/mm^2^ UV doses respectively. Scale bar is 100 µm. (B) The bar chart showing the live cell percentage 4h and 24h after the photopolymerization process. Error bars represent SEM and n = 4 for all data points. 71% and 76% viability were respectively measured 4h and 24h after the photopolymerization process at a UV dose of 20 mJ/mm^2^.

### Matrigel is a key component for obtaining stable BSPPs

In previous studies exploring the 3D culture of biliary epithelial cells^21,37^, type I collagen alone was insufficient to sustain the formation of any organized structure and Matrigel was required to form polarized structures with lumens^16,21,37^. Furthermore, the known association between biliary epithelial cells and basal lamina during bile duct morphogenesis led to the hypothesis that the basal lamina components are necessary to mimic the *in vivo* environment of developing bile ducts^21^. In order to evaluate this hypothesis, we supplemented the culture media, 2 days after photopolymerization, with 10% Matrigel whose composition closely mimics the components of the basal lamina, including especially laminin-1 that is crucial for epithelial polarization of mature cholangiocytes^21^. One week after photopolymerization, biliary epithelial cells in the samples without Matrigel failed to maintain organized structures **(Fig. 3, left panel)**, whereas in the presence of Matrigel, stable tubular and branched structures were maintained. The geometry of these structures correlated with the shapes on the original optical masks, with lengths up to 2 mm and widths ranging from 10 µm up to 100 µm for linear tubes **(Fig. 3, right panel and Fig. S2, day 7)**. Cells trapped within the proximal photopolymerization zones due to light scattering were also able to actively migrate and significantly remold the photopolymerized hydrogel structures between days 2 to 7. This resulted in modification of feature shapes and sizes compared to the ones dictated by original optical masks, especially when considering complex geometries such as smallest branches of trees (<20 µm). Seven days after photopolymerization, of which 5 days in culture media supplemented with Matrigel, we observed that the smallest branches of the 30° and 45° trees could not be resolved using our stereolithographic technique while the ones of 60° trees were often resolved (**Fig.S2, day 7**) and had cystic-shaped extremities.

**Figure 3.**
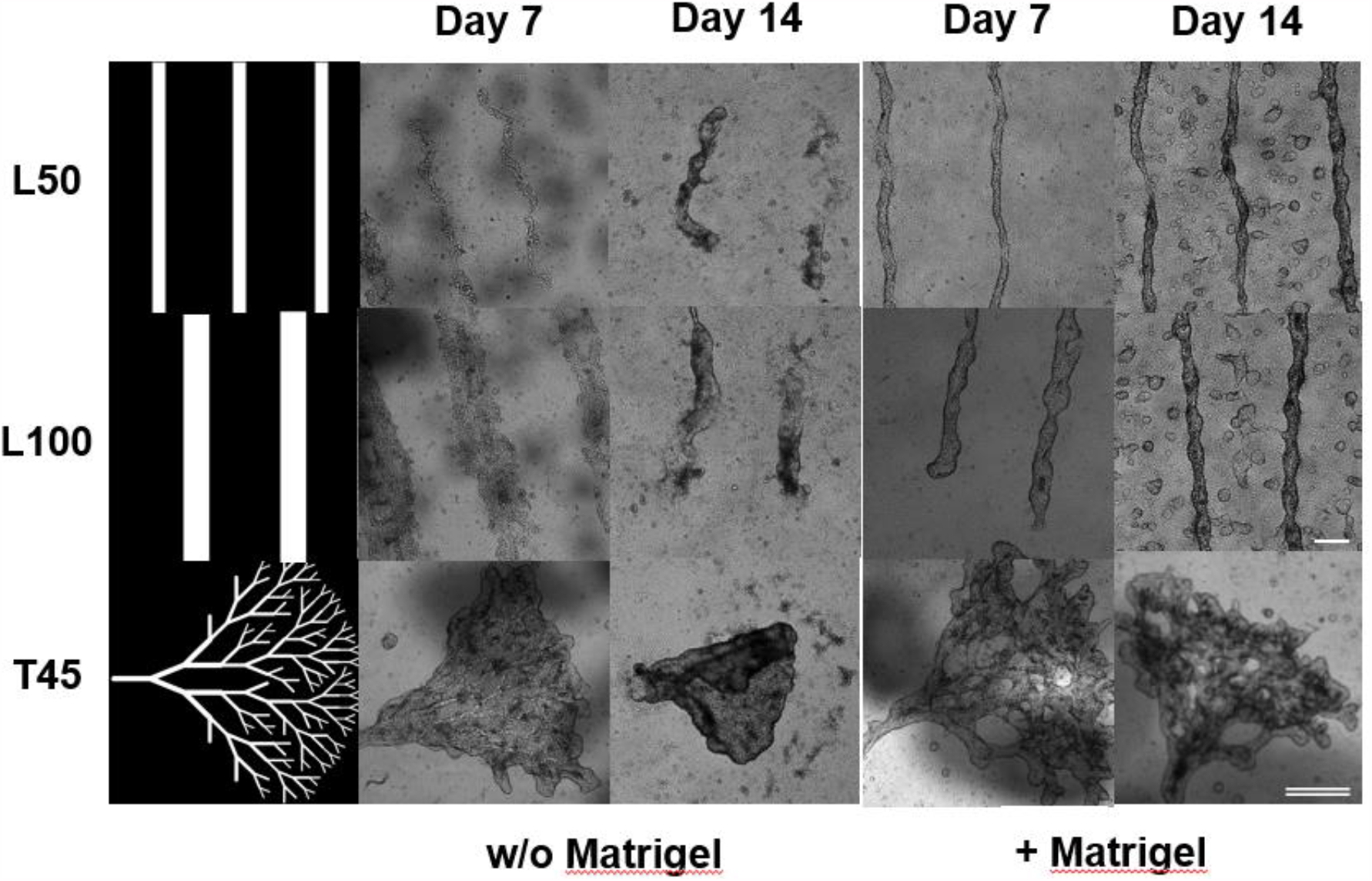
BSPP formation over time. Two days after fabrication and culture in cholangiocyte medium, BSPPs were divided into 2 groups and cultured respectively with and without Matrigel. Cholangiocytes were encapsulated either in 50 µm photopolymerized lines (L50), 100 µm photopolymerized lines (L100), and 45° photopolymerized trees (T45) and imaged after 7 and 14 days respectively. Single line scale bar is 200 µm (50-100 µm lines) and double-line scale bar is 1 mm (45° trees).

Two weeks after fabrication, in the samples without Matrigel, BSPPs were partially degraded and detached from the glass substrate (**Fig. 3 left panel, Fig.S3)** whereas, in the presence of Matrigel, BSPPs could be maintained up to 21 days **(Fig. 3 right panel, Fig. S2, day 14 and day 21, and Fig. S4)**.

These results suggest that DLP stereolithographic bioprinting can efficiently create a 3D ECM scaffold that guides proliferation, migration, and rearrangement of cholangiocytes when cultured in medium supplemented with Matrigel and that the resulting structures can be maintained for at least 21 days after photopolymerization.

**Figure S2.**
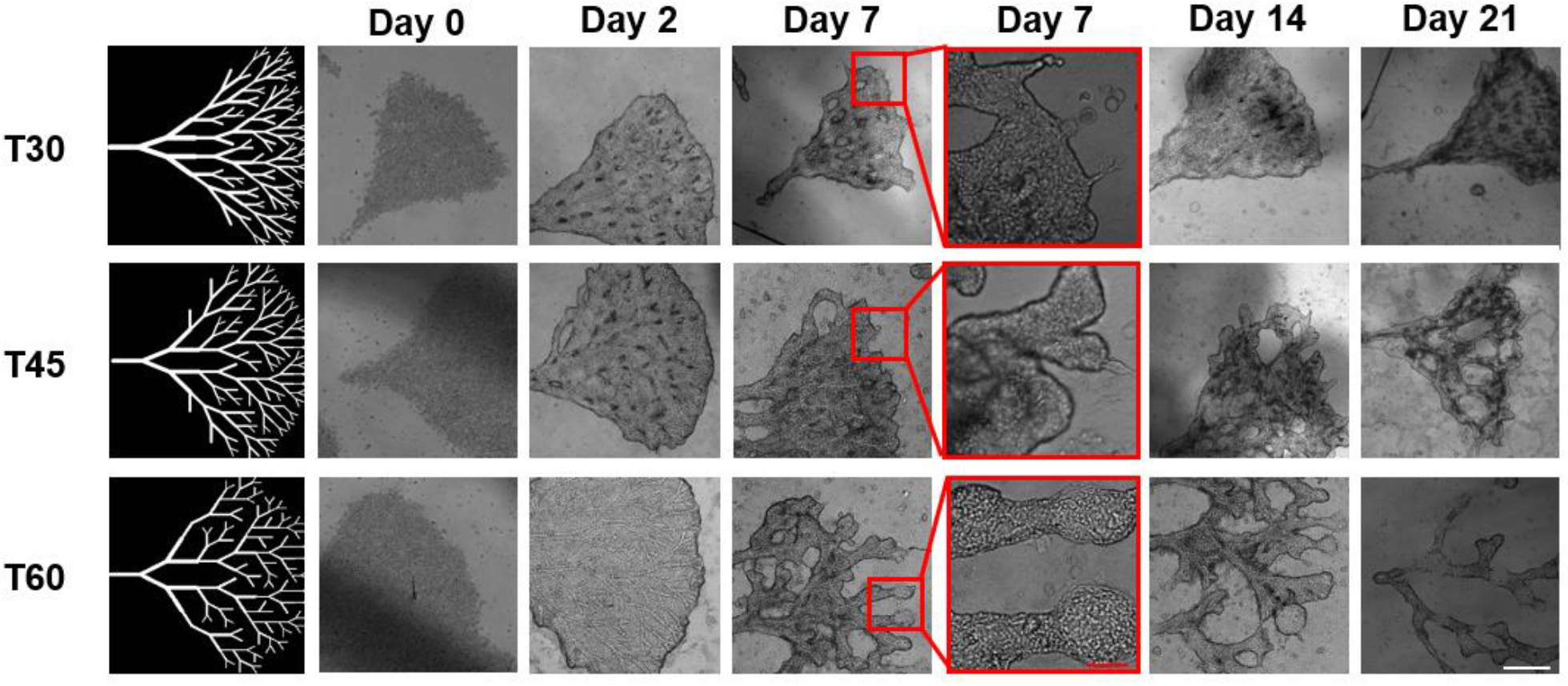
Cholangiocyte behavior in different tree geometries over 21 days of culture. Optical images showing the evolution of cholangiocyte structures after photopolymerization in a 30° tree (T30), a 45° tree (T45), and a 60° tree (T60) over time. Scale bar is 1 mm except for red-framed images where the red scale bar is 20 µm.

**Figure S3.**
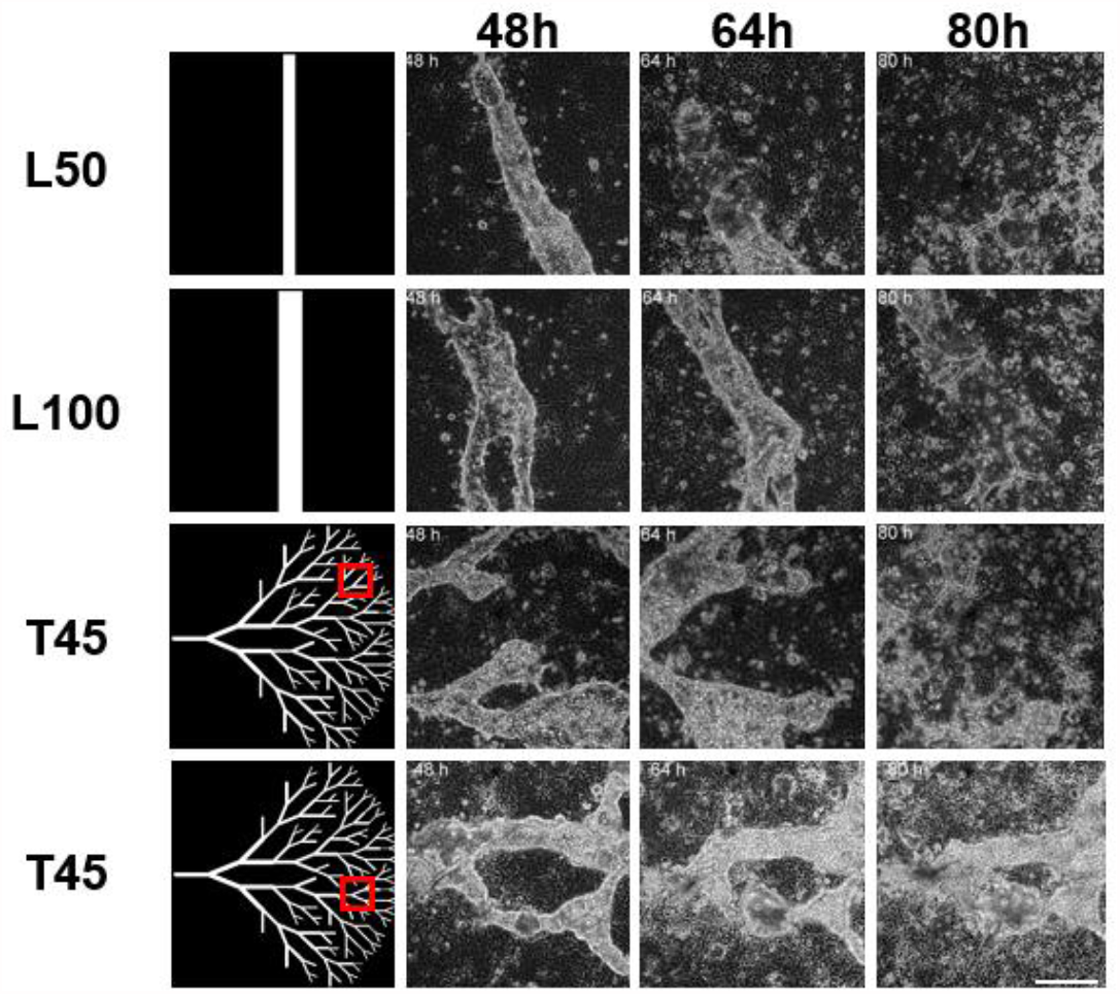
BSPPs cultured without Matrigel. Time-lapse transmission images of various photopolymerized structures encapsulating NRCs cultured without Matrigel. NRCs were encapsulated into 50 µm photopolymerized lines (L50), 100 µm photopolymerized lines (L100), and photopolymerized trees (T45). The image sequence shows the degradation of photopolymerized constructs over culture time at different time points: 48h, 64h, and 80h after fabrication. Scale bar is 100 µm.

**Figure S4.**
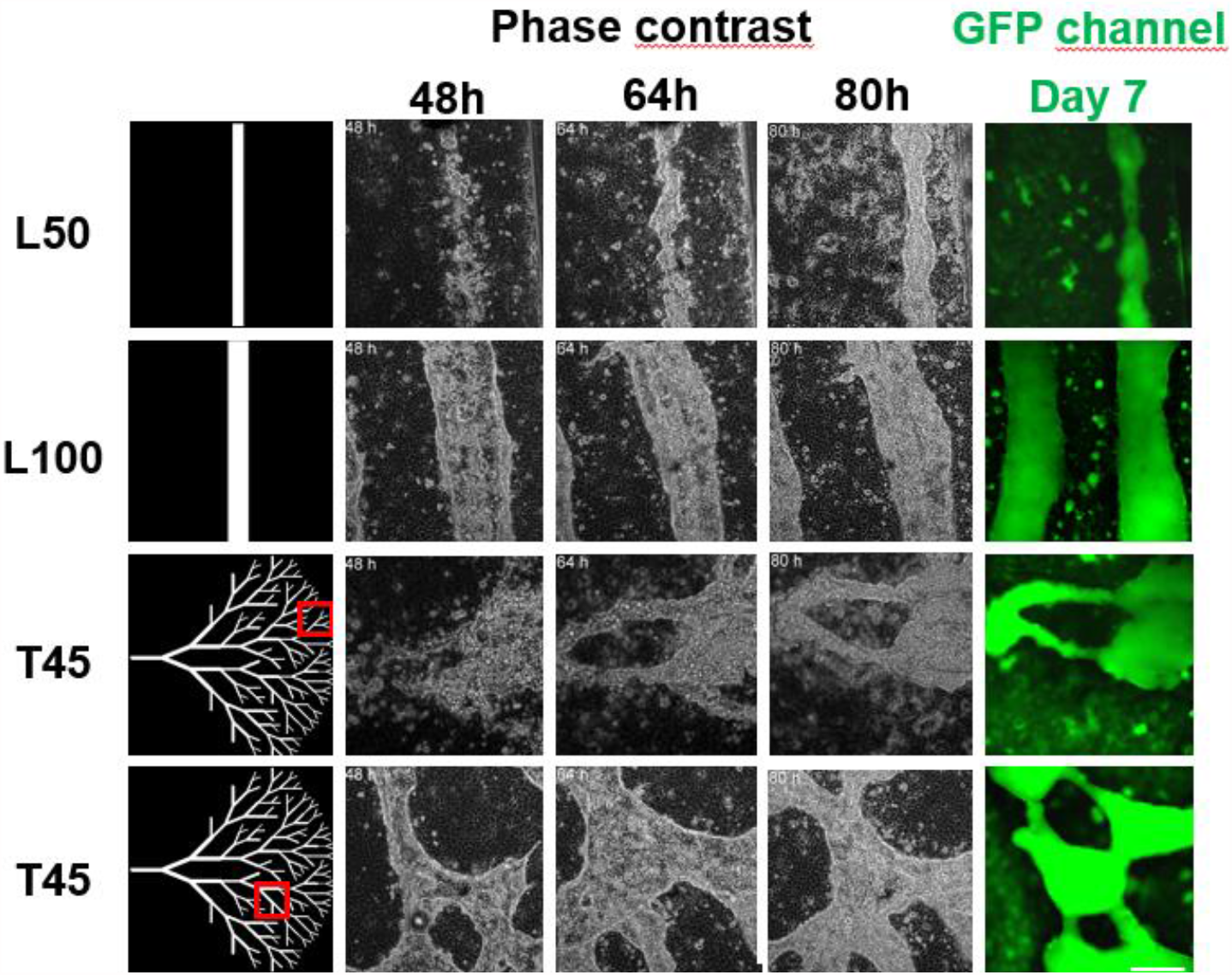
BSPPs cultured with Matrigel. Time-lapse transmission images of various BSPPs cultured with Matrigel. NRCs were encapsulated into 50 µm photopolymerized lines (L50), 100 µm photopolymerized lines (L100), and 45° photopolymerized trees (T45). Image sequences show the formation of biliary tubes over culture time at different time points: 48h, 64h, and 80h after fabrication. After 7 days of culture, for each implemented geometry, resulting BSPPs incubated with FDA show the accumulation of metabolized fluorescein within their luminal space. Scale bar is 100 µm.

### The obtained BSPPs have a monolayer wall and actively transport fluorescein from the cytoplasm into the lumen

Knowing that biliary epithelial cells spontaneously form cysts with a monolayer wall when cultured in natural or synthetic hydrogels^11,16^, we sought to investigate whether the BSPPs were forming lumens and whether these constructs could elicit the basic functions of a biliary epithelium.

After 7 days of culture, of which 5 days in medium supplemented with Matrigel, BSPPs were fixed, stained, and imaged using a laser scanning confocal microscope. Staining with phalloidin, anti-ZO1 antibodies, and DAPI revealed that the cholangiocytes in BSPPs are organized in a monolayer around a central lumen. These tubes could span hundreds of microns without any visible breaks in the walls. The diameter of the lumen was close to the size of the initial photopolymerized hydrogel structure as it can be seen in Fig.4: a tube with a 20-40 µm lumen **(Fig. 4A)** was formed on a 50 µm photopolymerized line, while the tube with the 10-15 µm lumen **(Fig. 4B)** resulted from a 20 µm photopolymerized line. Moreover, F-actin distribution was characteristic of epithelial cells with dense bands along cell-cell borders. Cholangiocytes formed tight junctions as shown by ZO-1 staining suggesting the formation of an intact epithelial barrier.

**Figure 4.**
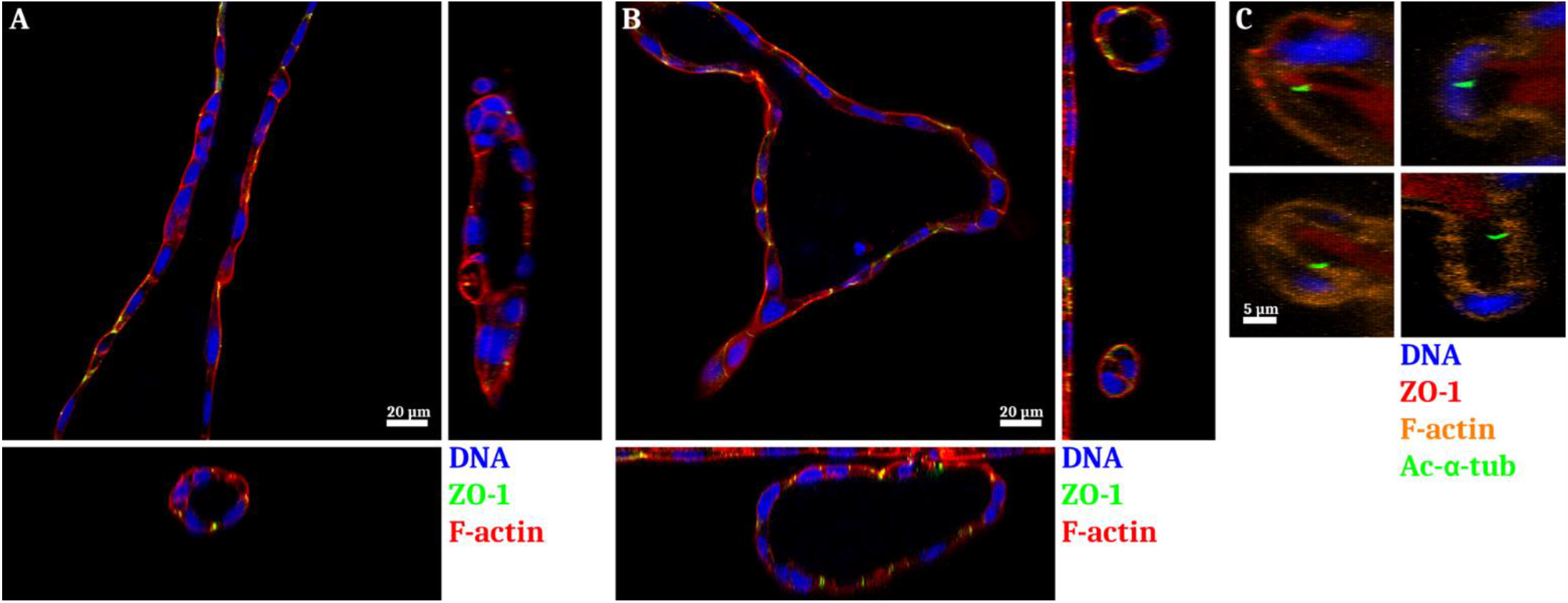
Confocal imaging of BSPPs after 7 days of culture in media supplemented with Matrigel for 5 days. NRC form single-layer tubular structures from an (A) 50 µm photopolymerized line and (B) 20 µm photopolymerized line. Staining for F-actin (phalloidin), tight junctions (anti-ZO-1 immunostaining), and DNA (DAPI). (C) Additional staining for acetylated α-tubulin reveals cilia facing the lumen of our tubular structures and formed by NRCs (orange – F-actin; red – ZO-1; green – acetylated α-tubulin; blue – DNA). Confocal imaging using Zeiss LSM 780 equipped with a 40x objective. Scale bar – 20 µm for A and B, 5 µm for C.

We further characterized the polarization of the cholangiocytes in BSPP tubes by staining them for acetylated alpha-tubulin - a component of the primary cilia. A number of cells had developed primary cilia and these were all facing the lumen (**Fig. 4C**).

Functional characterization of BSPP tubes was performed using the fluorescein diacetate (FDA) assay. FDA is a non-fluorescent molecule, which living cells hydrolyze to form fluorescent fluorescein in their cytoplasm. Cholangiocytes, unlike most non-epithelial cells can further secrete fluorescein through specific MRP2^38–41^ transporters located on their apical membranes^42–44^. As shown in Fig. 5A, our BSPPs actively transported fluorescein into the lumen, in both linear BSPPs of different diameters and branched BSPPs. The same result was obtained with Rhodamine 123 – a fluorescent compound secreted by MDR-1 – another anion transporter localized on the apical membrane (**Fig. S6**).

**Figure 5.**
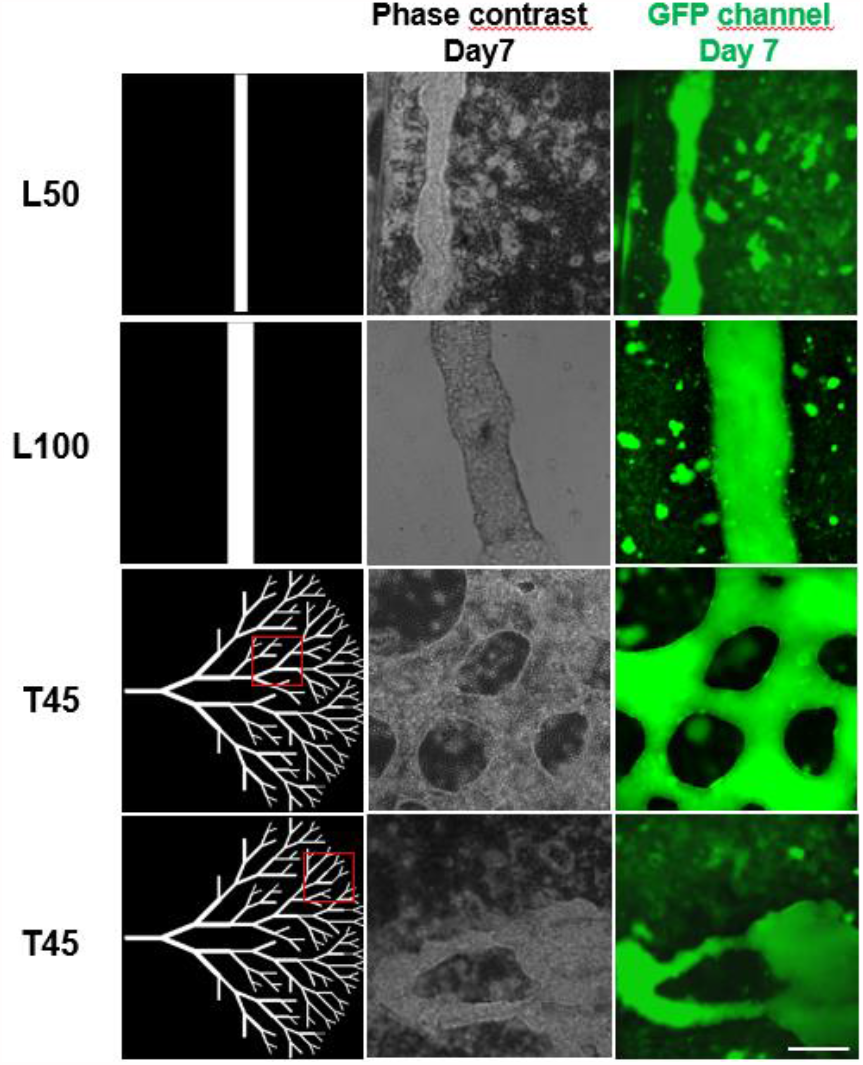
Functional characterization of BSPPs with a FDA assay. 7-day-old BSPPs were incubated with FDA and imaged with fluorescence microscope to evaluate the transport of fluorescein inside the lumen. Scale bar is 100 µm.

Taken together, these results demonstrate that we are able to produce geometrically controlled biliary tubes possessing structural and functional properties of intrahepatic bile ducts using DLP stereolithography to encapsulate cholangiocytes in soft hydrogels.

**Figure S5.**
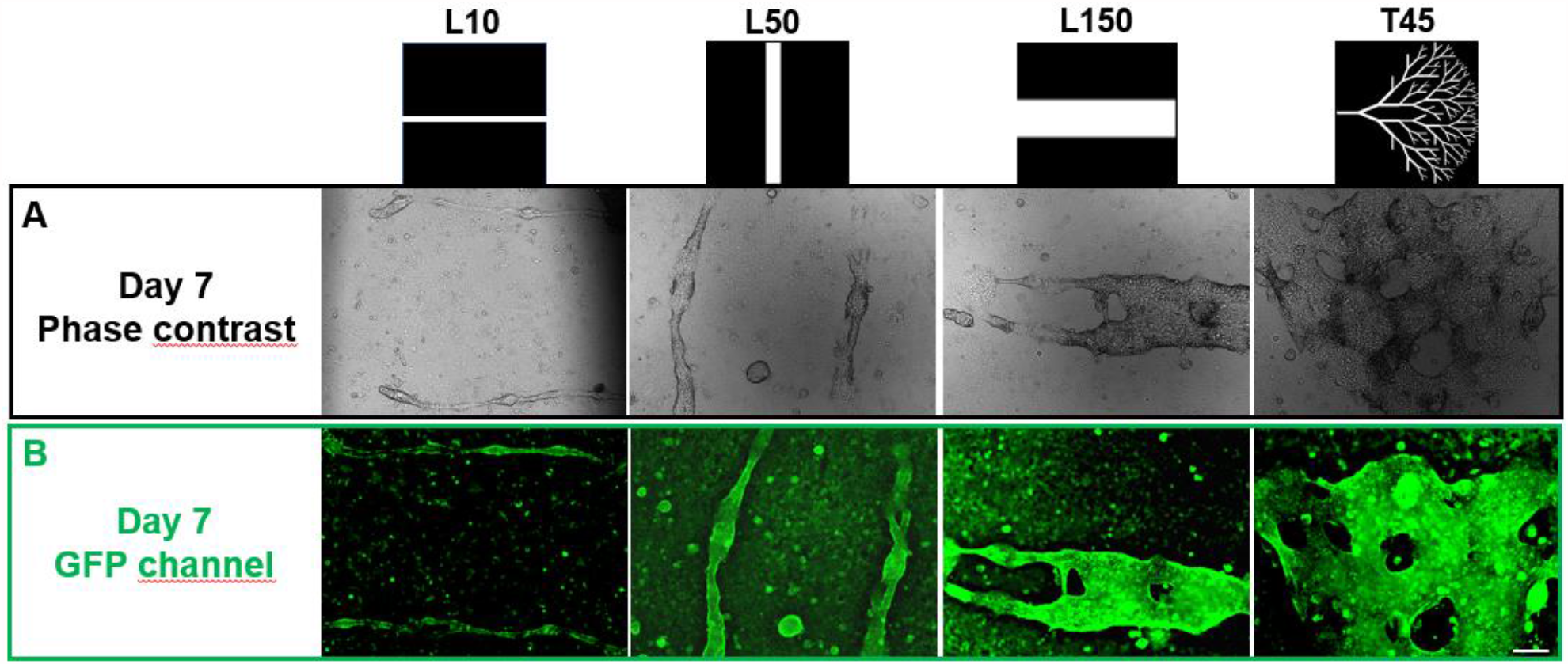
FDA assay using 21-day-old BSSPs. (A)Transmission and (B) fluorescent images of BSPPs after incubation with FDA. After 21 days of culture, resulting BSPPs incubated with FDA show the accumulation of metabolized fluorescein within their luminal space for each implemented geometry. Scale bar is 100 µm.

**Figure S6.**
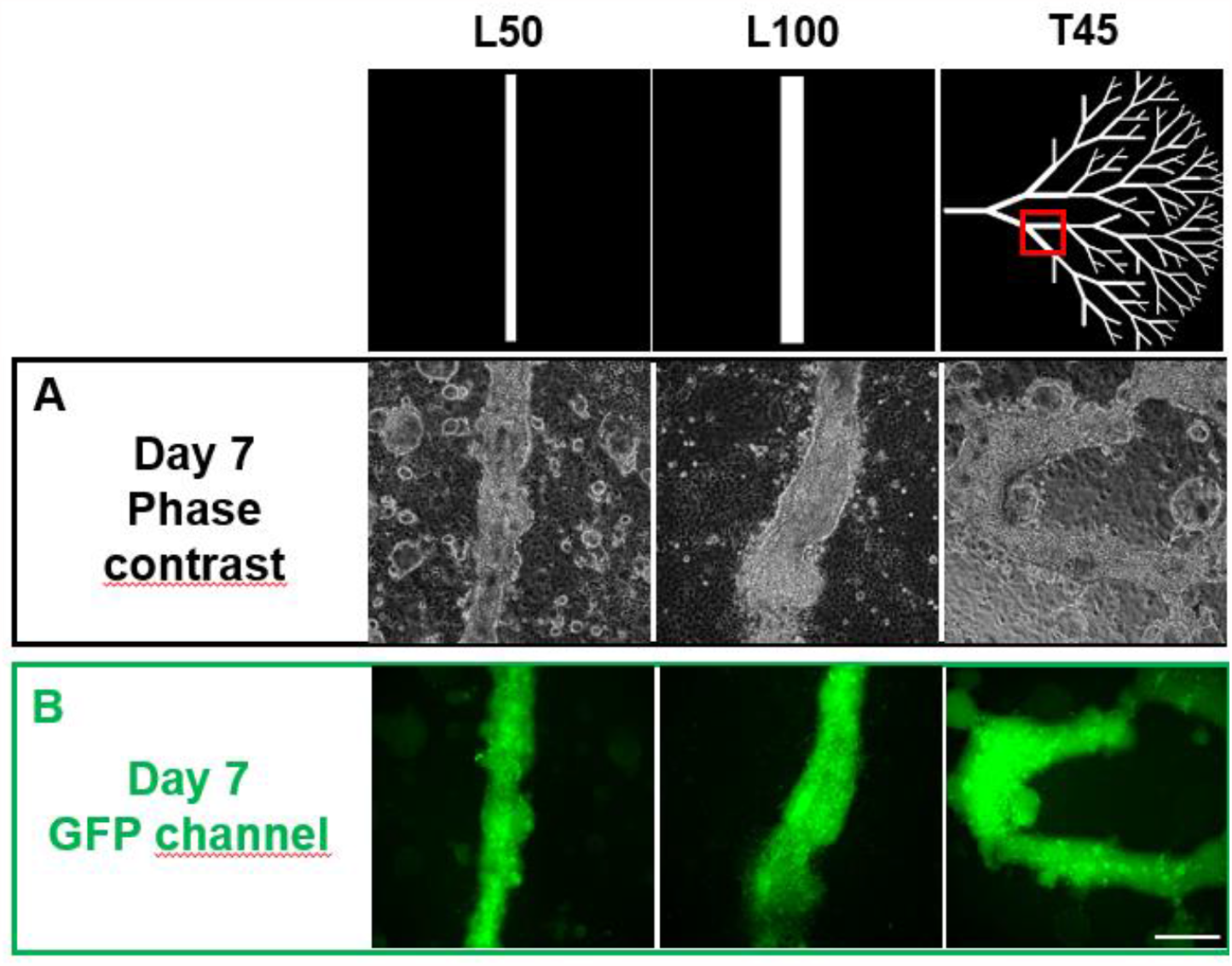
Rhodamine 123 assay using 7-day-old BSSPs. (A)Transmission and (B) corresponding fluorescent images of BSPPs after incubation with Rhodamine 123. Rhodamine 123 accumulated in the lumen of BSPPs reflecting the functional activity of NRC membrane channel multidrug resistance protein 1. Scale bar is 100 µm.

## Discussion

Cholangiocytes, like many epithelial cells, are known to spontaneously form cysts, sometimes adopting an elongated tubular structure^37,45^. However, the absence of control over their size and shape prevents the development of reproducible models of functional polarized intra-lobular biliary ducts. In this work, we demonstrate that the control over the initial geometry of a hydrogel by DLP stereolithography can allow the guided self-organization and polarization of embedded cholangiocytes into biliary tubes with predefined geometries including linear tubes with diameters found in the intrahepatic region of the biliary tree and, most remarkably, branched networks in tree-like configurations. Hence, this work stands as a striking illustration of how engineering techniques, in particular micropatterning as well as bioprinting, can help unleash the full potential of cellular self-organization into reproducibly building functional tissues and mini-organs with higher fidelity and robustness as discussed previously in our review^46^.

The composition of the gel was a critical parameter and the conjunction of all three types of matrices proved to be necessary to sustain self-organization. The precise role of each component has yet to be elucidated, although hypotheses can be made based on what is already known about these natural extracellular matrix polymers. Type I collagen is by far the most abundant protein in all vertebrates. It auto-assembles into heterofibrils with either type III or type V collagen. Heterofibrils are the structural and mechanical scaffold of most parenchymal tissues such as lung, kidney, liver, muscle, or spleen. Type I collagen is in particular present in normal livers in the blood vessel walls and the wall of biliary ducts of the portal tracts^47^. C^MA^ alone was rapidly degraded in the presence of cells in our initial experiments and the addition of HA^MA^ improved the durability of the resulting photopolymerized hydrogel. Indeed, hyaluronic acid is known to improve the mechanical properties of constructs in tissue engineering applications. It is also known to improve cell migration of cornea epithelial cells^48^ and both these properties could play a role in promoting efficient and stable tissue self-organization in this study. FG in turn plays a critical role in inflammation, wound healing, and angiogenesis by interacting with blood cells, endothelial cells, and other cell types when spreading into the extravascular space. FG has been implicated in particular as a substratum for epithelial cell migration during wound repair^49,50^. Even more interestingly, Guadiz et al. suggested that polarized secretion of FG by lung epithelium toward the epithelial basement membrane may behave as a substrate for cell adhesion and migration during wound healing^51^. In light of our results with BSPPs, FG was indeed essential for cholangiocyte tube formation as shown in Fig. 2 where NRC structures copying the initial photopattern are seen after 7 days with the C^MA^, HA^MA^, and FG mix and not with C^MA^ or HA^MA^ alone.

As illustrated by **Fig. 3** and the supplementary **movies S7** **and S8**, tubes form by proliferation and collective migration of cholangiocytes which seem to engulf the 3D structures, working their way between the glass and the hydrogel and eventually closing the tube when two sheets meet over the hydrogel. In some movies, in particular **movies S8B and S8D**, a “zipper” type of closing of these two sheets can be observed. These tubes form within a few days and Matrigel plays a critical role in the stability of these tubes over time: at early time points up to the third or fourth day after photopolymerization, depending on the initial geometry of the construct, functional tubes can be observed without Matrigel but these are unstable and collapse rapidly (**movies S7**). In the presence of Matrigel, tubes are stable, functional, and can be maintained for 21 days. At later time points, transport assays show that these tubes dissipated the fluorescent dye very quickly from the central lumen, pointing to defects in the overall integrity of the tube after 21 days (**Fig. S5**). The process whereby Matrigel stabilizes these tubes has yet to be elucidated. In our conditions, the concentration of Matrigel does not form a gel, which means that its role is not one of physical support. We hypothesize that a component of the ECM, most probably laminin, is reinforcing the polarization of the tube by offering a basal signal to the outside of the tube.

**Movies S7:**
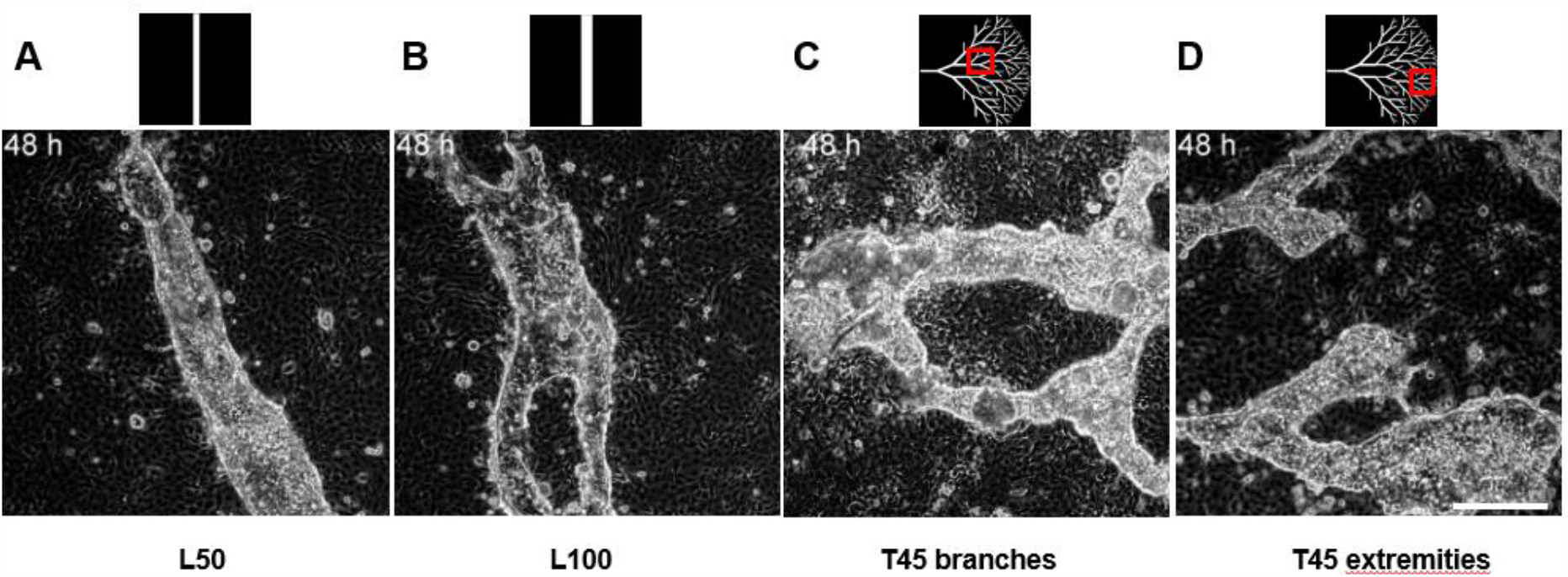
Timelapse recordings in the absence of Matrigel at day 2 - Two days after fabrication, BSPPs were cultured in medium without Matrigel. Cholangiocytes were imaged for 2 or 3 days either in (A) 50 µm photopolymerized lines, (B) 100 µm photopolymerized lines, and (C-D) 45° photopolymerized trees.

**Movies S8:**
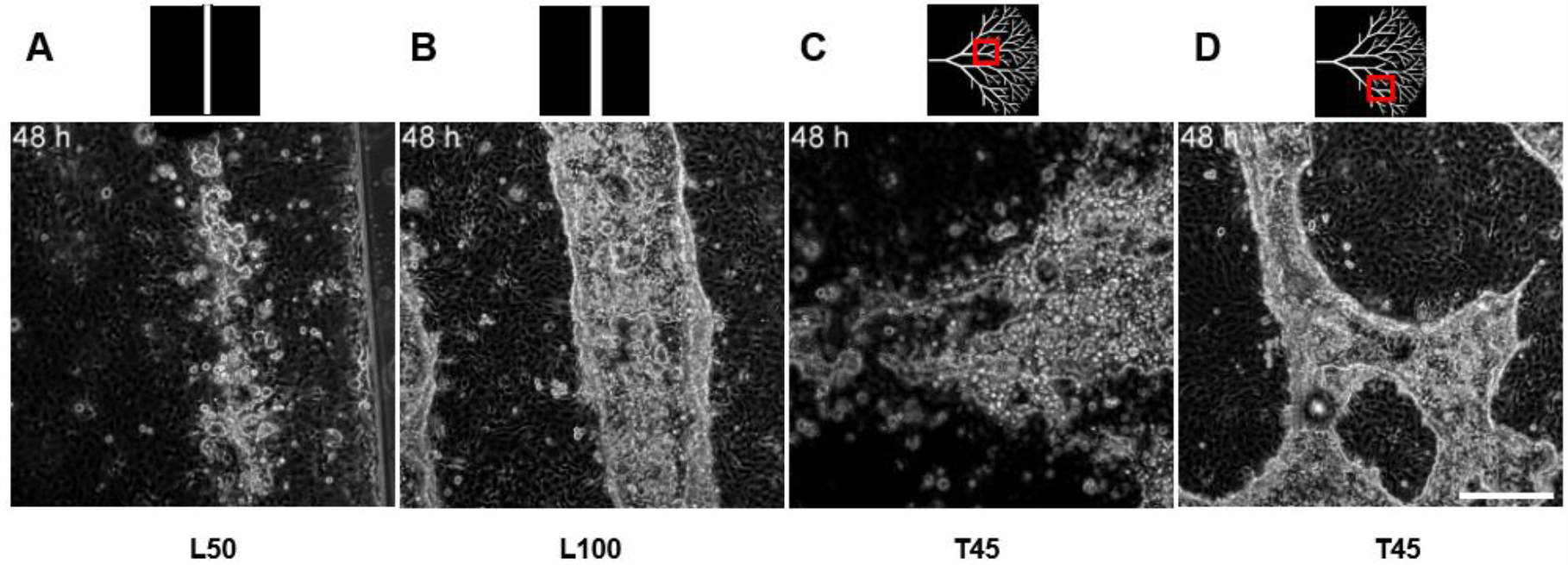
Timelapse recordings in the presence of Matrigel at day 2 - Two days after fabrication, BSPPs were cultured in medium with Matrigel. Cholangiocytes were imaged for 2 or 3 days either in (A) 50 µm photopolymerized lines, (B) 100 µm photopolymerized lines, and (C-D) 45° photopolymerized trees.

It is interesting to note that the collapse mechanism observed without Matrigel initiates from a central point and forms circular expanding structures that break up the initially condensed tubular structure. We hypothesize that the polarity of cells is randomly distributed in these constructs without the basal signaling brought by the laminin-containing Matrigel. It then appears that an avalanche effect leads to a collective collapse initiating from a central node. In itself, this mechanism would be interesting to elucidate further.

Finally, from an application standpoint, the dimensions of our BSPP tubes are consistent with those present in the liver lobule making them the first model of intrahepatic bile tube organogenesis which can prove invaluable to test receptor-mediated transport functions, drug transport in the liver and help elucidate precise mechanisms and gene functions during the early development of the biliary tree, as well as interaction with other cell types. In contrast to what is obtained inside bulk porous hydrogels, the tubes obtained in these conditions offer full access to the outside of the tube, allowing to introduce other cell types after BSPP maturation, such as hepatocytes and endothelial cells. Even more so, the tree-like design represents a planar mimic of the intrahepatic biliary network that collects bile into bigger bile ducts. Similarly, it is envisioned that the secreted bile in the BSPP could be collected and analyzed in an integrated microfluidic circuit opening new horizons in liver-on-chip development and applications.

In parallel, from a clinical research point of view, these structures have the added advantage of being produced in a thin single plane and cell sheet technology^52^ could be used to recover the structures and stack them with hepatocyte cell sheets to obtain more complex liver tissue constructs.

## Material and methods

### 1. Culture of cholangiocytes in 2D

NRC, a cholangiocyte cell line developed by Vroman and LaRusso^27^ were maintained on collagen-coated culture vessels in Dulbecco’s modified eagle medium F12 (DMEM F12, Gibco), non-essential amino acids (Gibco), lipid concentrate (Gibco), vitamin solution (Gibco), l-glutamine (Gibco), soybean trypsin inhibitor (Sigma), dexamethasone (Sigma), ethanolamine (Sigma), bovine pituitary extract (Gibco), Triiodo-L-thryronine (Sigma), Insulin-transferrin-Selenium (Gibco), and penicillin-streptomycin (Gibco) supplemented with 5% fetal bovine serum as described by De Groen et al.^53^

NRCs from passages 3-12 were used for the 3D stereolithography experiments.

### 2. Fabrication of BSPP using DLP stereolithography

Before 3D stereolithography, photopolymerizable hydrogels were prepared with 0.2% (w/v) type I methacrylated collagen (C^MA^, Advanced BioMatrix), 0.3% (w/v) methacrylated hyaluronic acid (HA^MA^, Advanced BioMatrix), 0.14% (w/v) fibrinogen (FG, Sigma), and 1% (w/v) LAP (Allevi). NRC were harvested with 0.25% trypsin-EDTA (Sigma) and diluted to cell concentrations of 10^7^ NRC/mL in photopolymerizable hydrogel.

The 3D stereolithography experiments were performed using an Olympus IX83-inverted microscope (Olympus Corporation) equipped with a DMD-based UV patterned illumination device (PRIMO, Alveole). The wavelength of the laser is 375 nm. Rasterized image files are sent to the DMD device which is an array of approximately two million micromirrors that can be controlled individually to generate the optical pattern. The optical photopattern is then projected through a microscope lens onto the photopolymerizable hydrogel containing NRCs.

Photopolymerizable hydrogels containing 10^7^ NRCs/mL were dispensed by 30 µL with a pipette into the space between a methacrylated glass coverslip and a polydimethylsiloxane (PDMS) stamp. The whole device was then loaded to the microscope slide holder and exposed to the UV pattern through a 10x at 20 mJ/mm^2^ doses. After UV exposure, the PDMS stamp was peeled off and unpolymerized part of photopolymerizable hydrogel containing NRC was removed and washed off with PBS buffer (Sigma).

### 3. Culture of BSPPs

After 3D stereolithography, the BSPPs were transferred to a petri-dish or a well plate and cultured in NRC culture medium. For the Matrigel-treated BSPPs, NRC culture medium was supplemented with 10% of Matrigel (Sigma) 2 days after photopolymerization. For all samples, culture media were renewed 3 times a week.

### 4. Fluorescent dye transport assay

On day 7 or day 21 of culture, BSPPs were incubated with fresh serum-free medium containing 15 μM of fluorescein diacetate (FDA, Sigma) for 5 min at 37°C and were then washed with serum-free medium three times. BSPPs were visualized using an Olympus IX83-inverted microscope with a 10x objective or Leica DMi8 microscope equipped with 10x objective and images were taken every 10 min for 1h. Temperature and CO2 concentration were kept at 37°C and 5%, respectively.

### 5. Fixation, immunofluorescent staining, and confocal microscopy of BSPP

To investigate the biliary epithelium formation *in vitro*, BSPPs cultured for a week in NRC culture medium supplemented with 10% of Matrigel for four days were fixed in 4% PFA in PBS for 15 min at room temperature, washed three times for 10 min in PBS and incubated for 1 hour in blocking solution (5% normal goat serum (Cell Signaling Technology #5425), 0,3% Triton X-100 in PBS) at room temperature. The blocking was followed by overnight incubation with the primary antibodies (1% BSA, 0,3% Triton X-100 in PBS) at +4°. After three 10 min washings in PBS at room temperature, samples were incubated with secondary antibodies (1% BSA, 0,3% Triton X-100 in PBS) for 2 hours at room temperature. Finally, samples were mounted using Aqua-Poly/Mount mounting medium and imaged using a confocal laser scanning microscope Zeiss LSM780 equipped with 40x objective. Below is the list of used antibodies and dyes with the dilution ratios relative to stock concentrations recommended by the manufacturers:

- anti-Acetyl-α-Tubulin (Cell Signaling Technology #5335) – 1:800;
- anti-ZO-1 (Invitrogen #33-9100) – 1:100;
- goat anti-Mouse IgG (Cy3) (Abcam #ab97035) – 1:200;
- goat anti-Rabbit IgG (Alexa Fluor 488) (Thermo Fisher Scientific #A-11034) – 1:200;
- phalloidin (Alexa Fluor 647) (Thermo Fisher Scientific #A22287) – 1:200;
- DAPI (Roche #10 236 276 001) – 4 µg/ml

### 6. Time-lapse image acquisition of BSPPs

Time-lapse microscopy was performed at 37°C and 5% of CO2, with images taken at 30-min intervals using an Olympus IX83-inverted microscope or a Leica DMi8 microscope equipped with a 10x objective.

### 7. Photopolymerizable hydrogel compositions

Before 3D stereolithography, various photopolymerizable hydrogel formulations were prepared: (i) C^MA^/HA^MA^/FG with 0.2% (w/v) C^MA^, 0.3 % (w/v) HA^MA^, 0.14% (w/v) FG, and 1% (w/v) LAP (ii) C^MA^/HA^MA^ with 0.2% (w/v) C^MA^, 0.3 % (w/v) HA^MA^, and 1% (w/v) LAP (iii) C^MA^/FG with 0.2% (w/v) C^MA^, 0.14 % (w/v) FG, and 1% (w/v) LAP (iv) HA^MA^/FG with 0.3 % (w/v) HA^MA^, 0.14 % (w/v) FG, and 1% (w/v) LAP (v) 0.2% (w/v) C^MA^ and 1% (w/v) LAP (vi) HA^MA^ with 0.3 % (w/v) HA^MA^ and 1% (w/v) LAP (vii) FG with 0.14 % (w/v) FG and 1% (w/v) LAP. Each previously described photopolymerizable hydrogel mix was gently mixed with NRCs at a cell concentration of 10^7^ NRCs/mL and were photopolymerized with a 20 mJ/mm^2^ dose. After being incubated in NRC medium for 48 h at 37°C, the photopolymerized samples were cultured during 5 more days in NRC culture medium supplemented with 10% Matrigel.

### 8. Rheological analysis

Rheological features of photopolymerized hydrogels were measured by a Discovery HR 2 rheometer (TA Instruments) equipped with a parallel plate geometry (d= 25 mm). Two different photopolymerized hydrogel formulations were prepared: C^MA^/HA^MA^/FG (0.2% C^MA^, 0.3 % HA^MA^ 0.14% FG, 1% LAP) and C^MA^/HA^MA^ (0.2% C^MA^, 0.3 % HA^MA^, 1% LAP). Samples with a thickness of 100 µm and 25 mm in diameter were photopolymerized with a 20 mJ/mm^2^ dose. After being incubated in NRC medium for 24 h at 37°C, the photopolymerized samples were subjected to oscillatory measurements. In order to eliminate the environmental noise in data acquisition, the frequency sweeps were conducted at room temperature in the range of 0.1-10 Hz, and with a constant strain rate of 1%, considered to be in the linear viscoelastic range.

## Funding and acknowledgments

This work received the financial support of the iLite RHU program (grant ANR ANR-16-RHUS-0005).

We thank Didier Letourneur and Isabelle Bataille (INSERM U1148, Univ Paris 13) for access to their rheology equipment. We also thank Latifa Bouzhir for her assistance with NRC culture. We finally thank Benoit Vianay from the CytoMorphoLab (INSERM, CEA, U976) for his technical assistance with the DLP stereolithography setup.

## Author contributions

E.M-A. D.A. F.C. and A.F. conceived and designed the experiments; E.M-A. D.A., E.G. and W.F. performed the experimental work and contributed to the manipulation of materials, reagents, and analysis tools; E.M-A. D.A. W.F., P.D-W, F.C., J.L and A.F. analyzed the data and wrote the manuscript. All authors read and approved the final manuscript.

## Data availability

The raw/processed data required to reproduce these findings cannot be shared on a permanent website at this time due to technical or time limitations. The raw/processed data required to reproduce these findings can be shared by the authors upon request.

